# A Novel Role for the Nuclear Receptor, NR4A1, in *Klebsiella pneumoniae* Lung Infection

**DOI:** 10.1101/2020.09.03.282475

**Authors:** Jessica Partyka, Matthew Henkel, Brian T. Campfield

## Abstract

*Klebsiella pneumoniae* is a Gram-negative bacterial pathogen and common cause of pneumonia and bacteremia. Increasingly, *K. pneumoniae* has become a public health concern due to its rate of nosocomial infection and emerging, broad-spectrum antibiotic resistance. The nuclear receptor NR4A1 exhibits functionality in a multitude of organ systems and is implicated as having a role in the immune response to bacterial infection, though its role in *K. pneumoniae* infection is unknown. To determine if *Nr4a1* functions in response to *K. pneumoniae* pulmonary disease, we infected wild-type and *Nr4a1*^*−/−*^ mice with *K. pneumoniae* and assessed bacterial growth, immune cell recruitment and function, and cytokine production. We found that *Nr4a1*^*−/−*^ mice had increased bacterial burden in the lungs and spleen, though no differences in cell recruitment. Pro-inflammatory cytokines, *Il1β* and *Il6*, as well as chemokine, *Cxcl2*, were significantly decreased in the BAL fluid cells of *Nr4a1*^*−/−*^ mice 5 hours post-infection. Additionally, *Nr4a1*^*−/−*^ mice had reduced IL-1β and myeloperoxidase protein production. We then examined the bactericidal function of macrophages and neutrophils from WT and *Nr4a1*^*−/−*^ mice. We identified that *Nr4a1*^*−/−*^ neutrophils had decreased bactericidal function compared to wild-type neutrophils, which was associated with reduced expression of *Il1β*, *Lcn2*, *Mpo*, and *Lyz2*. These data suggest *Nr4a1* plays a novel and essential role in neutrophil function during the host immune response to *K. pneumoniae* pulmonary infection.

## INTRODUCTION

*Klebsiella pneumoniae* is a Gram-negative bacterium of the *Enterobacteriaceae* family. Recognized as an intestinal commensal in healthy individuals, *K. pneumoniae* is a common cause of disease in immunocompromised patients and is the third most common healthcare associated infection, manifesting as urinary tract infections, bacteremia, and pneumonia (1). In addition, *K. pneumoniae* strains are becoming increasingly resistant to antibiotic treatments, including carbapenems and extended-spectrum β-lactams (ESBL); thus, *K. pneumoniae* constitutes a public health threat for which new treatment modalities are urgently needed (2, 3).

The orphan nuclear receptor subfamily 4 group A member 1 (NR4A1) is a steroid-thyroid receptor and intracellular transcription factor encoded by the *Nr4a1* gene, also known as Nur77. NR4A1 is a molecule with diverse biologic functions, including regulation of inflammation in the central nervous system, heart, and lung (4–11). Specifically, NR4A1 is a factor in macrophage development and inflammatory response through polarization and transcriptional regulation (12–16). Within macrophages, NR4A1 exhibits a direct effect on the NF-κB pathway by directly blocking p65 from binding to the κB element, and prior work has shown deletion of the *Nr4a1* gene increases p65 phosphorylation, thus activating NF-κB mediated transcription, in macrophages (17, 18).

Due to its varied impact in immunological pathways, the role of NR4A1 in bacterial infection has been explored but remains poorly understood. In *Escherichia coli* induced peritonitis, NR4A1 has a limited role in the clearance of the bacteria (19). However, Hamers *et al.* later showed the deletion of the *Nr4a1* gene aggravated *E. coli* induced peritonitis due to pro-inflammatory macrophage polarization (20). In addition to showing NR4A1 directly interacts with the NF-κB pathway, Li *et al.* also showed NR4A1 is a vital protein for inflammation reduction and, ultimately, survival in a sepsis model (17). The sole study of NR4A1 in bacterial lung infection examined *E. coli*, an uncommon cause of pneumonia, and showed that *Nr4a1* was rapidly induced during infection, and that *Nr4a1*^*−/−*^ mice had decreased bacterial burden *in vivo*, though the mechanisms underlying these observations remain unclear (21).

The current study examined the role of NR4A1 in *K. pneumoniae* pneumonia. Here, we found that *Nr4a1* deficiency led to an increase of *K. pneumoniae* bacterial lung burden and dissemination. Mechanistically, phagocyte recruitment was unaffected, though inflammatory cytokine/chemokine production was decreased in *Nr4a1*^*−/−*^ mice, identifying that *Nr4a1*^*−/−*^ neutrophils had defective bactericidal function and attenuated inflammatory gene expression, suggesting that *Nr4a1* is a novel, critical mediator of neutrophil function.

## RESULTS

### *Nr4a1*^*−/−*^ mice are more susceptible to *Klebsiella pneumoniae* pulmonary infection

To explore the impact of *Nr4a1* on control of *K. pneumoniae* pulmonary infection, WT and *Nr4a1*^*−/−*^ mice had bacterial burden determined by colony-forming units (CFU) at 24 and 48 hours after infection in the lung, as a measure of local control, and in the spleen as a measure of bacterial dissemination. Compared to WT mice, *Nr4a1*^*−/−*^ mice had significantly increased *K. pneumoniae* lung burden and spleen burden 48 hours post-infection (Fig. 1c-d). These differences were not present at 24 hours post-infection (Fig. 1a-b). These data identify that *Nr4a1*^*−/−*^ mice controlled *K. pneumoniae* acute lung infection poorly, suggesting that *Nr4a1* may play an important role in the immune response to acute *K. pneumoniae* lung infection.

**Figure 1.**
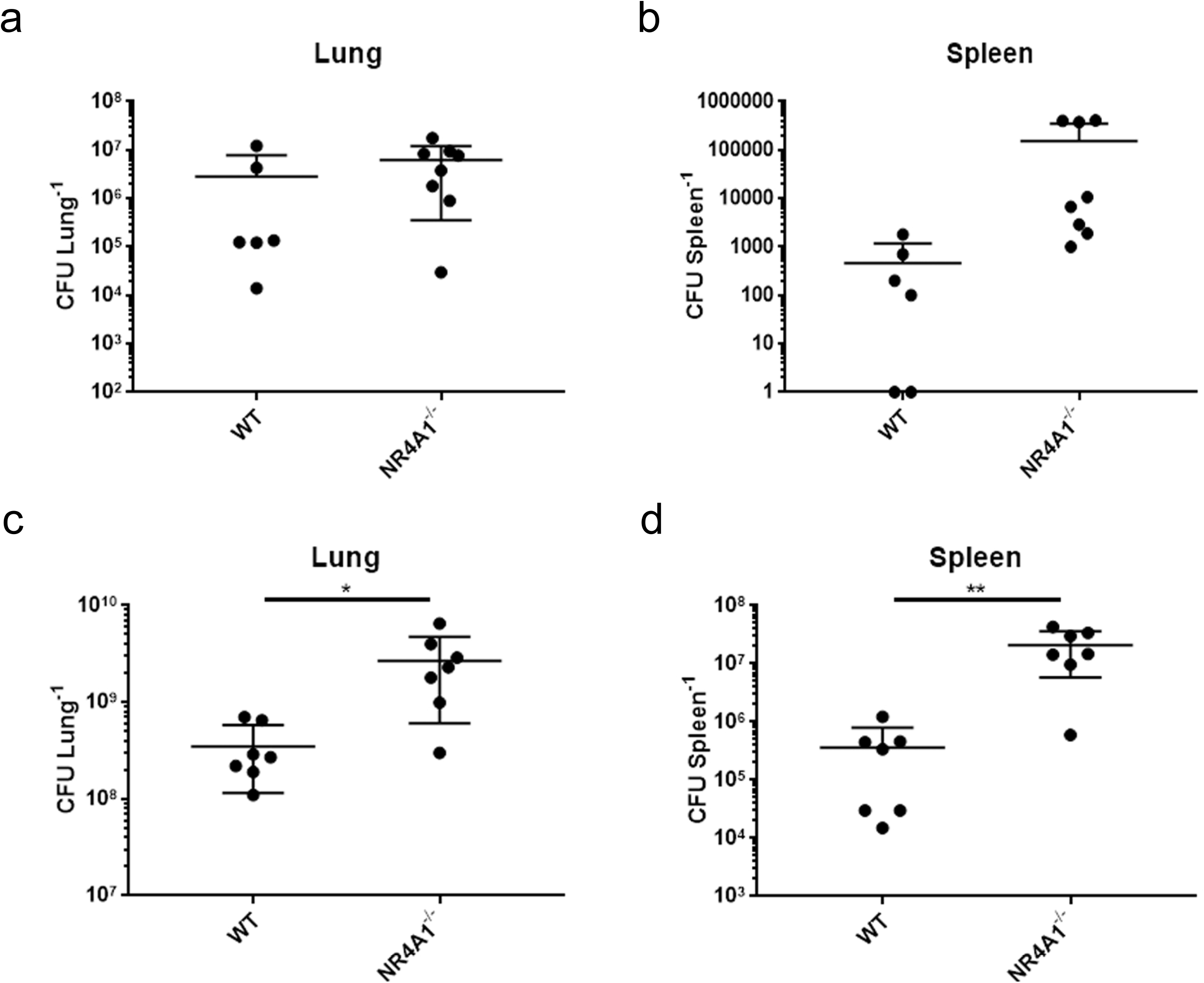
*Nr4a1*^*−/−*^ mice are more susceptible to *Klebsiella pneumoniae* pulmonary infection. The **(a)** lung burden and **(b)** spleen burden of Wild-type (WT, *n*=6) and *Nr4a1* knockout (*Nr4a1*^*−/−*^, n=8) mice 24 hours post pulmonary *K. pneumoniae* infection. The **(c)** *K. pneumoniae* lung burden and **(d)** spleen burden in WT (*n*=7) and *Nr4a1*^*−/−*^ (*n*=7) mice 48 hours post pulmonary infection. Panels **(a)** and **(b**) for the 24-hour lung and spleen burdens are representative of two experiments. Panels **(c**) and **(d**) for the 48-hour lung and spleen burdens are pooled from two experiments. Significance was determined by the Student’s *t*-test. *\*P* < 0.05, *\*\*P* < 0.01

### *Nr4a1*^*−/−*^ mice have reduced pro-inflammatory cytokine production in BAL fluid cells and whole lung early following *K. pneumoniae* lung infection

Next, to determine if immediate phagocyte function was impaired in the absence of *Nr4a1*, we analyzed cytokine/chemokine gene expression and protein production in the bronchoalveolar lavage fluid (BALF) and lung from WT and *Nr4a1*^*−/−*^ mice 5 hours post *K. pneumoniae* pulmonary infection. Pro-inflammatory cytokines, *Il1β* and *Il6*, and the neutrophil chemokine, *Cxcl2*, were significantly decreased in the BALF of *Nr4a1*^*−/−*^ mice (Fig. 2a, b, e). However, *Tnfa* and the chemokines *Cxcl1* and *Cxcl5* were not different (Fig. 2c, d, f), suggesting the NR4A1-dependent gene regulation was reflective of a specific cell population or pathway mediating the phenotype and this was observable within 5 hours post-infection.

**Figure 2.**
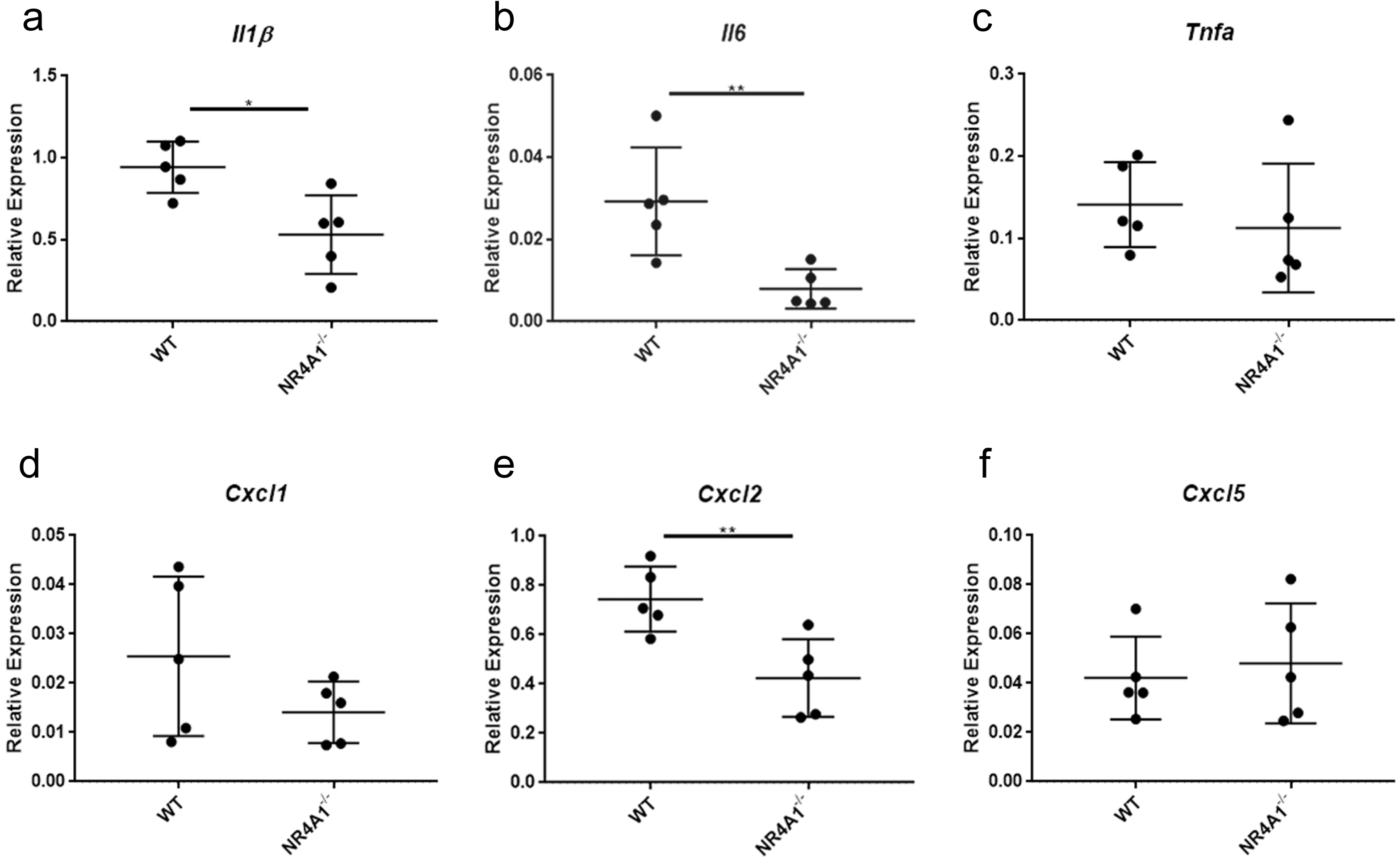
*Nr4a1*^*−/−*^ mice have selectively reduced transcription of pro-inflammatory cytokines and chemokines in BAL cells early post-infection. Transcript abundance for pro-inflammatory cytokines **(a)** *Il1β*, **(b)** *Il6*, and **(c)** *Tnfa*, as well as chemokines **(d)** *Cxcl1*, **(e)** *Cxcl2*, and (f) *Cxcl5* from the BAL fluid cells of WT (*n*=5) and *Nr4a1*^*−/−*^ (*n*=5) mice 5 hours after *K. pneumoniae* infection. These data are representative of two experiments. All gene transcripts were compared to the housekeeper beta-2-microglobulin (*B2M*). Significance was determined by the Student’s *t*-test. *\*P* < 0.05, ** *P* < 0.01

Whole lung protein from WT and *Nr4a1*^*−/−*^ mice was also assessed at 5 hours and 24 hours post *K. pneumoniae* infection. Consistent with early gene expression, IL-1β protein was significantly reduced in *Nr4a1*^*−/−*^ compared to WT (Fig. 3a) at 5 hours post-infection. This reduction in IL-1β protein was sustained through 24 hours post-infection (Fig. 3c). We then sought to determine whether the Nr4a1-dependent phenotype was associated with reduced production of the effector molecule myeloperoxidase (MPO), an enzyme critical for phagocyte-mediated bacterial killing of *K. pneumoniae* (22). In the *Nr4a1*^*−/−*^ strain, MPO production trended down within 5 hours post-infection (*P* = 0.0689) (Fig. 3b). By the 24 hours post-infection timepoint, *Nr4a1*^*−/−*^ mice had significantly decreased MPO protein in the lungs compared to WT mice (Fig. 3d). These data suggest that early immune gene transcription reflects protein production during *K. pneumoniae* lung infection, and that *Nr4a1*-deficient susceptibility to infection is associated with reduced production of phagocyte markers.

**Figure 3.**
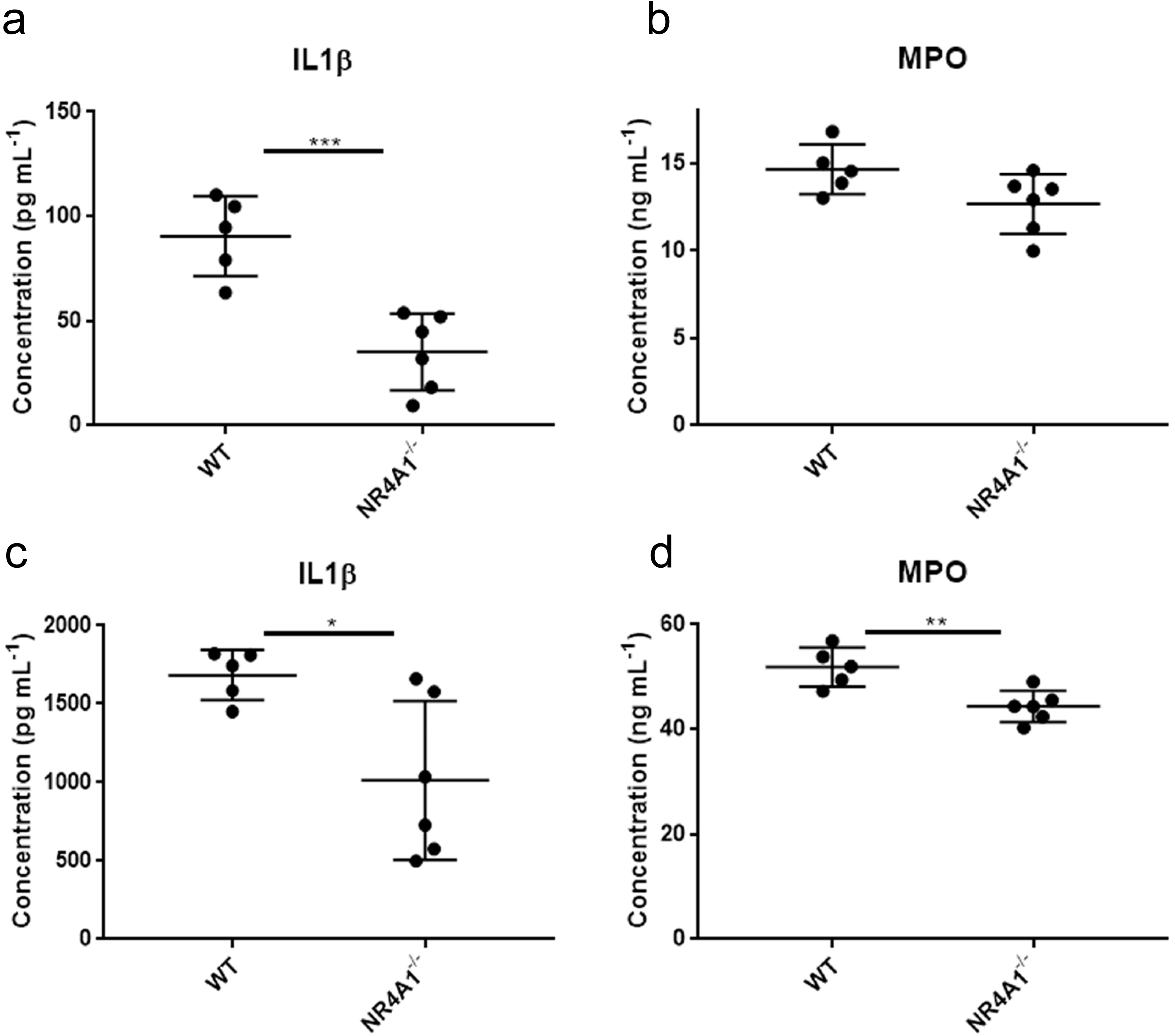
*Nr4a1*^*−/−*^ mice have reduced IL1β and MPO lung protein after *K. pneumoniae* lung infection. *Nr4a1*^*−/−*^ mice (*n*=6) have reduced **(a)** IL1β 5 hours post-infection with *K. pneumoniae* compared to WT controls (*n*=5), though **(b)** MPO is not significantly reduced. By 24 hours post infection, *Nr4a1*^*−/−*^ mice (*n*=6) have significantly reduced **(c)** IL1β and **(d)** MPO protein compared to WT mice controls (*n*=5). These data are all representative of two experiments. Significance was determined by the Student’s *t*-test. \**P* < 0.05, \*\**P* < 0.01, \*\*\**P* < 0.005

### *Nr4a1*^*−/−*^ and WT mice have similar immune cell abundance and cytokine gene expression 24 hours following *K. pneumoniae* lung infection

To examine a potential mechanism by which *Nr4a1* protects against lung infection, we performed flow cytometry on the lungs of WT and *Nr4a1*^*−/−*^ mice at 24 hours post *K. pneumoniae* infection, prior to differences in bacterial burden, to determine if *Nr4a1* deletion resulted in a change in immune cell recruitment. No significant differences were observed in inflammatory macrophages (CD45^+^/CD11b^+^/CD11c^+^), alveolar macrophages (CD45^+^/CD11c^+^/SiglecF^+^), or neutrophils (CD45^+^/CD11b^+^/Ly6G^+^) (Fig. S1b-d). Interestingly, no differences in transcript were found between pro-inflammatory cytokines *Il1β, Tnfa*, or *Il6*, or the neutrophil-specific gene *Lcn2* (Fig. S2a-d). These data suggest the role *Nr4a1* plays in the immune response to *K. pneumoniae* acute lung infection is not due to cell recruitment and is not reflected in gene transcription through 24 hours post-infection.

### *Nr4a1*^*−/−*^ macrophage and neutrophil bactericidal function *in vitro*

Bone marrow macrophage bacteria killing assays were analyzed to determine bacterial growth as a measure of *in vitro* bactericidal function from WT and *Nr4a1*^*−/−*^ mice. No differences in bacterial growth were observed between the two strains at 30 minutes or 60 minutes co-culture with *K. pneumoniae* (Fig. S4a), identifying that macrophages of both genotypes have similar bactericidal function. Similarly, the gene transcript levels for *Il1β*, *Il6*, and *Cxcl2* were not different, suggesting that the differences seen in the BALF cells and lung homogenate *in vivo* does not likely reflect *Nr4a1*-mediated function in macrophages (Fig. S4b-d).

Neutrophil bacteria killing assays were similarly analyzed to determine bacterial growth as a measure of *in vitro* bactericidal function from WT and *Nr4a1*^*−/−*^ mice. Notably, in WT neutrophils co-cultured with *K. pneumoniae, Nr4a1* gene transcription is induced compared to naïve WT neutrophils (Fig. 4a). Supernatants from *Nr4a1*^*−/−*^ neutrophils had significantly higher *K. pneumoniae* growth compared to WT neutrophils within 60 minutes, though at 30 minutes no differences were observed between genotypes (Fig. 4b-c). These data suggest that *Nr4a1*^*−/−*^ neutrophils *K. pneumoniae* killing is time dependent. Taken together, these data suggest *Nr4a1* transcription is induced in neutrophils by *K. pneumoniae* infection and is critical for the bactericidal activity.

**Figure 4.**
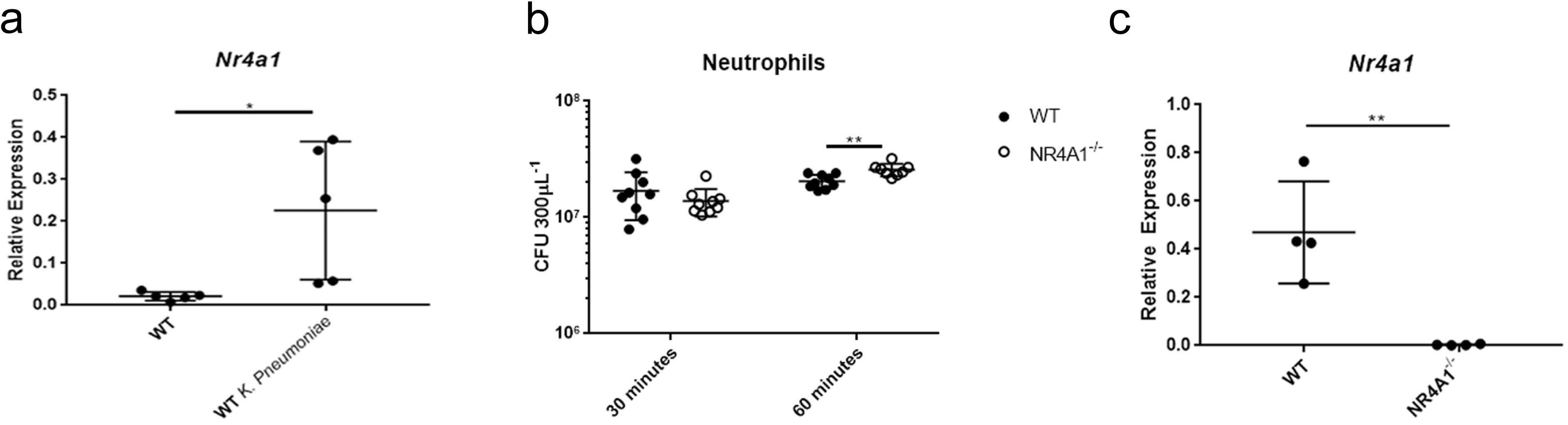
*Nr4a1* is essential for neutrophil anti-bactericidal function. **(a)** *Nr4a1* gene expression of unstimulated (*n*=5) and *K. pneumoniae* co-cultured (*n*=5) WT neutrophils. *Nr4a1* gene expression is significantly increased in WT neutrophils co-cultured with *K. pneumoniae*. The **(b)** *K. pneumoniae* burden from the supernatants of WT (*n*=9) and *Nr4a1*^*−/−*^ (*n*=9) neutrophils at 30 minutes and 60 minutes (WT *n*=9, *Nr4a1*^*−/−*^ n=8) after co-culture with *K. pneumoniae*. **(c)** *Nr4a1* gene expression of WT (*n*=5) and *Nr4a1*^*−/−*^ (*n*=5) neutrophils 60 minutes after *K. pneumoniae* co-culture. WT neutrophils have significantly more *Nr4a1* gene transcript than *Nr4a1*^*−/−*^ neutrophils. All gene transcripts were normalized to *B2M*. The burden results in panel **(b)** are pooled from two separate experiments and representative of five experiments total. Panels **(a)** and **(c)** are representative of three experiments. Significance was determined by the Student’s *t*-test. *\*P* < 0.05, *\*\*P* < 0.01, *\*\*\*\*P* < 0.0001

### *Nr4a1*^*−/−*^ neutrophils have decreased cytokine transcript compared to WT neutrophils

To assess *Nr4a1*-dependent transcriptional changes in neutrophils, *Nr4a1*^*−/−*^ neutrophil RNA was isolated 30 minutes after *K. pneumoniae* co-culture (when the bacterial burdens are equivalent). Here, *Nr4a1*^*−/−*^ neutrophils had decreased *Il1β* transcript compared to WT neutrophils, as was seen in BALF cells 5 hours post-infection (Fig. 5a). *Nr4a1*^*−/−*^ neutrophils also had significantly decreased expression of neutrophil antibacterial transcripts *Lcn2*, *Mpo*, and *Lyz2* (Fig. 5b, c, d), further suggesting an important role for *Nr4a1* in neutrophil response to *K. pneumoniae* infection.

**Figure 5.**
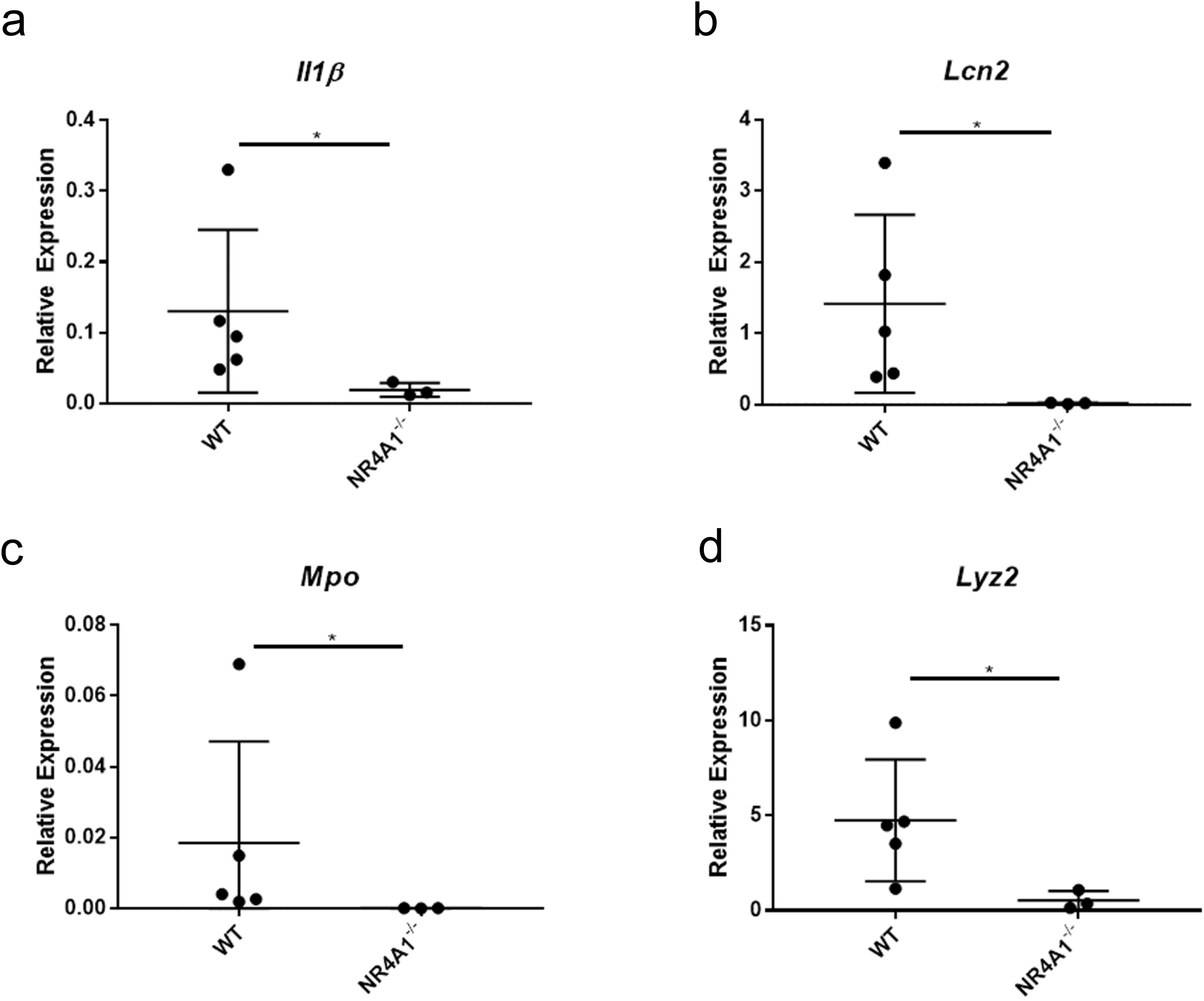
*Nr4a1*^*−/−*^ neutrophils have decreased inflammatory gene expression following *K. pneumoniae* infection. *Nr4a1*^*−/−*^ neutrophils (*n*=3) have decreased genetic expression of **(a**) *Il1β* and neutrophil-specific genes **(b)** *Lcn2*, **(c)** *Mpo*, and **(d)** *Lyz2* compared to WT controls (*n=*5). These data are all representative of five experiments. All gene transcripts were normalized to *B2M*. Significance was determined by the Mann-Whitney *U*-test. *\*P* < 0.05

## DISCUSSION

Due to its nosocomial infection rate and developing antibiotic resistance, *Klebsiella pneumoniae* is a growing public health concern (1–3). Developing new approaches to treat *K. pneumoniae* and other antibiotic resistant infections is an increasingly urgent task. One of the methods through which this may be done is targeting the host immune response to stimulate better clearance of bacterial burden from the infection site. This can be done by targeting different immune cells, including T-cells, macrophages, and neutrophils, to enhance response to infection (23–25). For macrophages and neutrophils, this can be done by augmenting phagocytosis of bacteria (24–27). However, modulating host immunity is an understudied approach for the treatment of bacterial infection. More study of host immune responses and host-pathogen interaction in bacterial infection, including *K. pneumoniae*, is required if harnessing host immunity as a potential treatment is to be successful.

The orphan nuclear receptor NR4A1 is expressed in many immune cells, such as T-cells and myeloid cells. NR4A1 is necessary for proper T-cell function and development (28, 29). It has likewise been shown to be important for the development and function of several subsets of macrophages, and as an important regulator of macrophage polarization (12–16). Little is known about the role of NR4A1 in neutrophils, though human and murine neutrophils express all three of the transcripts for Nur nuclear receptors, *Nr4a1*/Nur77, *Nr4a2*/Nurr1, and *Nr4a3*/Nor1, in response to inflammatory stimuli (30, 31). Highly expressed in numerous immune cell types, we sought to identify cellular mechanisms for NR4A1 in order to elucidate its potential as a target for immunomodulatory therapies.

The current study aimed to determine the role of NR4A1 in *K. pneumoniae* pulmonary infection. Prior studies have described a limited or detrimental role for NR4A1 in bacterial infections (19, 21). To date, the only study of bacterial lung infection employed an *E. coli* model, in which *Nr4a1*^*−/−*^ mice had improved bacterial clearance and survival (21). However, the clinical relevance is limited, as *E. coli* is not a cause of community or hospital acquired pneumonia, and little mechanistic information was ascertained from this work. Conversely, our study determined that *Nr4a1* loss was detrimental in the *K. pneumoniae* pneumonia model. Mechanistically, the observation that NR4A1 deficiency did not impair macrophage or neutrophil cell recruitment to the lung suggested that NR4A1 may be important for innate antibacterial cell-intrinsic function. While some genes were unchanged, reduced lung expression of the innate pro-inflammatory cytokines, *Il1β* and *Il6,* as well as the neutrophil recruitment chemokine, *Cxcl2*, 5 hours after infection suggested an innate phagocyte response pathway defect (32). Several disease models have shown that macrophage or monocyte expression of NR4A1 has significant impact on inflammation in models of cardiovascular disease, LPS-induced sepsis, and fibrosis (6, 7, 17, 18, 33, 34). However, in each of these models, NR4A1 modulated macrophage inflammatory genes in response to insult or the deletion of *Nr4a1* resulted in an *exaggerated* macrophage inflammatory phenotype, while we observed *reduced* pro-inflammatory cytokines in *K. pneumoniae* infected *Nr4a1*^*−/−*^ lung tissue, which prompted further study into cell-specific responses to *K. pneumoniae* infection in the context of NR4A1.

Given that our work and others indicated *Nr4a1*^*−/−*^ macrophages had a dysregulated tolerization phenotype, it was unexpected that *Nr4a1*^*−/−*^ macrophages retained *K. pneumoniae* killing ability (18). Despite genetic profiles noting that *Nr4a1* is induced in neutrophils in response to inflammatory stimuli, no prior work has determined what role *Nr4a1* has in the neutrophil response, and the finding that *Nr4a1* is critically important in neutrophils, and not macrophages, has not been previously reported (30, 31). Specifically, reduced *Nr4a1*^*−/−*^ neutrophil *Mpo*, *Lcn2*, *Lyz2*, and *Il1β* gene expression suggests that *Nr4a1* deletion impairs antimicrobial responses by regulating cell-intrinsic early inflammatory gene expression. Alternatively, it may be that deletion of *Nr4a1* leads to a dysfunctional metabolic response in neutrophils following *K. pneumoniae* infection, as *Nr4a1* has profound impacts on T-cell and macrophage immunometabolism in several disease models (28, 35, 36). *Nr4a1*, as well as its family members *Nr4a2* and *Nr4a3*, are also known to be important in the stimulation of lipolysis and glucose utilization, and all are induced in neutrophils among other lipolysis-related genes in response to inflammatory stimuli (31, 37). Regardless, further study is necessary to dissect the molecular, cell-specific mechanism of NR4A1.

This work is not without limitations. Most prominently, all data represented here were performed in mouse models, and thus remains to be determined what role *Nr4a1* possesses in human neutrophil function. Though prior work suggests that *Nr4a1* is induced in human neutrophils responding to *E. coli*, human and murine neutrophils differ in numerous ways, including abundance and protein makeup (30, 38). Further study is required to discern what role, if any, NR4A1 has in the inflammatory response of human neutrophils. In addition, confirming the role of *Nr4a1* neutrophil-specific function *in vivo* remains undetermined and requires experimental approaches not currently available. We must also acknowledge that *in vitro* study of neutrophils is limited by the short, hours long half-life the cells have *ex vivo* without manipulation, complicating classical *in vitro* molecular and genetic experimental approaches (39).

Further study into the use of NR4A1 as a potential therapeutic target for host-immunity responses is also warranted. While the present study shows the deletion of *Nr4a1* to be detrimental to murine clearance of *K. pneumoniae* pneumonia through loss of neutrophil function, further study of the ability to rescue or enhance NR4A1 function is arranged in order to determine this potential, including the use of the NR4A1 agonist, cytosporone B, or the NR4A1 ligand-binding chemical, PDNPA, in the context of *K. pneumoniae* (11, 17, 40).

The current study also only looked at the role of NR4A1 in *K. pneumoniae* pneumonia. While *K. pneumoniae* is a prominent cause of pneumonia, it is not the only bacteria to do so, nor does *K. pneumoniae* only infect the lungs. More work should be done to determine if NR4A1 has a role in the response to other infectious bacteria (e.g. *S. pneumoniae*, *H. influenzae*) and viruses (coronavirus, influenza). Similarly, future studies should also be conducted to determine if the role of NR4A1 in host immune response is specific to the lungs, or if NR4A1 immune responses are important for other organ-specific infections.

The current study illustrates that *Nr4a1* is an important component in host immune response to *K. pneumoniae* pulmonary infection, specifically in neutrophils, marking NR4A1 as a potential target for future immunotherapies. Finding potential immunotherapy targets against bacterial infections is an increasing concern for the public health community as antibiotic-resistant strains continue to increase. Demonstrating the role NR4A1 has in host immune response to *K. pneumoniae* provides a rational biologic target around which to develop immunotherapies. Additionally, the novel finding that *Nr4a1* is an essential factor in neutrophil-specific response to *K. pneumoniae* infection can lead to future understanding of neutrophil-specific mechanisms in response to infection and how to modulate them.

## METHODS

### *Klebsiella pneumoniae* strain and inoculation

*Klebsiella pneumoniae* strain 396, a K1 serotype, was used for all experiments with a desired inoculation of 5×10^3^ colony forming units (CFU) per mouse for *in vivo* experiments and 2.5×10^6^ CFU per well for *in vitro* experiments. *K. pneumoniae* isolates were stored at −80°C in Luria-Bertani broth (LB; Thermo Fisher Scientific, Waltham, MA) supplemented with 15% glycerol.

Working cultures were grown overnight (18 hours) in 2mL of Trypticase soy broth (TSB; Sigma Aldrich, St. Louis, MO) at 37°C and 250 RPM. The next day, 20μL *K. pneumoniae* was sub-cultured into 2mL TSB and grown for ~2 hours at 37°C and 250 RPM to reach the bacterial growth log phase and a concentration of 1E9 CFU mL^−1^. The sub-cultured *K. pneumoniae* was then spun down at 5000G for 5 minutes and resuspended in 1mL 1XPBS (Gibco, Gaithersburg, MD). Sub-cultured *K. pneumoniae* was then diluted 1:10000 (1×10^5^ concentration) in 1XPBS for *in vivo* experiments or 1:20 (5×10^7^ concentration) for *in vitro* experiments. 50μL of 1×10^5^ inoculum was administered intratracheally to mice under 1:1 isoflurane for *in vivo* experiments. For *in vitro* experiments, 5mL of the 1:20 dilution was added to 5mL 1XPBS and 100μL of that solution was added to the well. Inoculum concentrations were confirmed by dilution series and LB agar plate CFU calculation.

### Animal models

Wild-type C57BL/6 mice (Jackson Laboratory, Bar Harbor, ME) and *Nr4a1* global knockout mice (*Nr4a1*^tm1Jmi^/J; Jackson Laboratory, Bar Harbor, ME) were housed in a pathogen-free environment with free access to autoclaved water and irradiated pellet food. All experiments were performed in accordance with the Institutional Animal Care and Use Committee (IACUC) of the University of Pittsburgh School of Medicine.

All *in vivo* experiments were performed on sex-matched mice between 8-10 weeks old. At the start of the experiment, mice were weighed and inoculated with 5×10^3^ *Klebsiella pneumoniae* CFUs. After 24 or 48 hours, mice were weighed again and euthanized under isoflurane and a secondary terminal bleed. The left lobe of the lung and the spleen was extracted into 1mL 1XPBS and homogenized for serial dilution and LB agar plate CFU calculation. The middle lobe of the right lung was extracted and placed into RNAlater (Invitrogen by Thermo Fisher Scientific, Carlsbad, CA) for RNA purification and the top and bottom lobes of the right lung were harvested for flow cytometry.

For BALF, 9-week-old male mice were inoculated with 5×10^3^ *Klebsiella pneumoniae* CFUs. After 5 hours, the mice were euthanized under isoflurane and a secondary terminal bleed and BALF was acquired by flushing the lungs with 1mL sterile 1XPBS. The 5-hour timepoint was selected based on prior LPS-induced acute lung injury (ALI) data suggesting neutrophil influx to the lungs starts as early as 2 hours after insult and as early as 3 hours after a gram-negative bacterial insult (41–43).

### Flow cytometry

To obtain single lung cell suspensions, lung lobe was collected and digested in DMEM containing 4mg mL^−1^ Collagenase (Sigma Aldrich, St. Louis, MO) and 0.2mg mL^−1^ Dnase (Sigma Aldrich, St. Louis, MO) at 37°C with agitation for 1 hour, followed by straining digest through a 70μm filter. Strained cell pellets were collected by centrifugation at 500xg for 5 minutes and red blood cells were lysed using ACK Lysing Buffer (Gibco, Carlsbad, CA) according to manufacturer recommendations. Total lung cells were enumerated with 0.2% trypan blue solution and an Invitrogen Countess automated cell counter.

Cells were then subjected to Fc blockade with anti-CD16/CD32 (Thermo Fisher Scientific, Waltham, MA), washed with 1XPBS, then stained with surface and/or intracellular flow cytometry antibodies specific to antigens of interest using eBioscience™ Foxp3 / Transcription Factor Staining Buffer Set (Thermo Fisher Scientific, Waltham, MA) according to manufacturer protocol. Cells were labeled for detection with surface antibodies to CD11c (HL3, BD Biosciences, San Jose, CA), SiglecF (E50-2440, BD Biosciences, San Jose, CA), CD11b (M1/70, Fisher Scientific, Waltham, MA) and Ly6G (1A8, BioLegend, San Diego, CA). Data acquisition was performed on an LSR Fortessa (SORP, BD Biosciences, San Jose, CA) using FACSDiva software version 8.0.1. Data was analyzed using FlowJo software version 10.1 (TreeStar, Ashland, OR). Cells were enumerated with 0.2% trypan blue solution and an Invitrogen Countess automated cell counter. Absolute cell numbers per mouse lung were enumerated using the total lung cell digest count and the flow cytometry percentage of their respective cell type.

### Bone marrow macrophage and neutrophil isolation

#### Macrophages

Age matched mice between 8-10 weeks old were euthanized under isoflurane and a secondary terminal bleed. Both femurs of each mouse were extracted and flushed with DMEM media

(Gibco, Carlsbad, CA) supplemented with 10% fetal bovine serum (FBS) and Penicillin-Streptomycin-Glutamine through a 100μm filter into a 50mL conical tube. Cells were spun down at 500xg for 5 minutes, red blood cell lysed with ACK lysis buffer (Gibco, Carlsbad, CA) for 1 minute, then spun down again at 500g for 5 minutes.

Again, cells were enumerated with 0.2% trypan blue solution and an Invitrogen Countess automated cell counter. 1×10^7^ cells were plated in 10cm dishes in 10mL of DMEM media supplemented with 10% FBS, Penicillin-Streptomycin-Glutamine, and M-CSF (Life Technologies, Carlsbad, CA) brought to a concentration of 20ng mL^−1^. Cells were incubated at 37°C and 5.0% CO_2_ for 7 days before being harvested in unsupplemented DMEM and used in the bacterial killing assay.

#### Neutrophils

Age matched mice between 8-10 weeks old were euthanized under isoflurane and a secondary terminal bleed. The isolation protocol was modified based on the neutrophil isolation protocol developed by Swamydas and Lionakis (44). Both femurs of each mouse were extracted and flushed with RPMI 1640 media (Fisher Scientific, Waltham, MA) supplemented with 10% FBS and 1mM EDTA (Gibco, Carlsbad, CA) through a 100μm filter into a 50mL conical tube. Cells were centrifuged for 7 minutes at 427xg and 4°C. Cells were then red blood cell lysed by adding 2mL 0.2% NaCl for 20 seconds and 2mL 1.6% NaCl. Cells were spun down again at the previous settings, washed with the supplemented RPMI media, and spun again. Cells were resuspended in 3mL PBS and counted using 0.2% trypan blue solution and an Invitrogen Countess automated cell counter.

Once counted, cells were overlaid on a 40%/70% Percoll (Fisher Scientific, Waltham, MA) gradient and centrifuged for 30 minutes at 427xg and 28°C. After 30 minutes, neutrophils were collected from the interface of the 40% and 70% Percoll layers, washed twice with unsupplemented RPMI 1640 and spun at 427xg and 4°C for 7 minutes, and resuspended in 1mL PBS. Neutrophils were counted and then plated in a 96-well plate at 2.5×10^5^ neutrophils/well. Neutrophils were incubated at 37°C and 5.0% CO_2_ for a 1-hour recovery period before being used in a bacterial killing assay.

### Bacterial killing assay

Both bone marrow macrophages and bone marrow neutrophils were individually plated in a 96-well plate at 2.5×10^5^ cells per well in their respective media. *K. pneumoniae* was added at a concentration of 2.5×10^6^ CFU per well for a multiplicity of infection (MOI) of 10. The plates were then incubated at 37°C and 5.0% CO_2_ on an orbital shaker set to 250 RPM.

Supernatants were collected at 30 and 60 minutes (for neutrophils) or 30 and 60 minutes (for macrophages) and the bacterial burdens of each well were calculated using serial dilutions of the supernatants and LB agar plate CFU calculations. In addition, cells were lysed using RNA lysis buffer (Invitrogen by Thermo Fisher Scientific, Carlsbad, CA) and stored for RNA extraction and qPCR analysis.

### RNA extraction and qPCR

For lung gene expression, the middle lobe of the right lung lobe was homogenized in 1mL of RNA lysis buffer (Invitrogen by Thermo Fisher Scientific, Carlsbad, CA) and RNA was purified per the manufacturer’s procedures. Cellular RNA was extracted by aspirating media and plating 300μL of the same RNA lysis buffer onto the cells directly after the killing assay. RNA was purified per the manufacturer’s procedures. RNA was quantified using the Nanodrop 2000 (Thermo Fisher Scientific, Waltham, MA) and 40ng of RNA for lung or 5-10ng of RNA for cells was converted to cDNA with iScript (Biorad, Hercules, CA). Real-time PCR was done on the cDNA with SYBR green master mix (Biorad, Hercules, CA) or Taqman master mix (Applied Biosystems, Foster City, CA) depending on primer probe. Primer probe assays were from ordered from Applied Biosciences or IDT. All lung gene expression was compared to the housekeeper gene hypoxanthine-guanine phosphoribosyltransferase (HPRT) and all cellular gene expression was compared to the housekeeper gene beta-2-microglobulin (B2M).

### ELISA protein analysis

For protein analysis of the whole lung, the left lung lobe was homogenized in 1mL of PBS and then diluted 1:10 to use in the IL1-β (Thermo Fisher Scientific, Waltham, MA) and MPO (R&D Systems, Minneapolis, MN) ELISAs. After collection of BALF, the BALF was spun at 500xg for 5 minutes to separate lung cells from the fluid. Undiluted BALF was used in the IL1-β and MPO ELISAs. All ELISAs were conducted per the manufacturer’s procedures. Plates were read at 450nm with a correctional wavelength of 540nm (Synergy H1, BioTek, Winooski, VT).

#### Statistical analysis

Investigators were not blinded to treatment but were blinded to individual/group during data analysis. All *in vivo* and *in vitro* statistical analyses were performed using Prism 7 (GraphPad, San Diego, CA). Briefly, all data are presented with mean ± SEM. All studies comparing two groups were analyzed by two-sided student’s *t*-test or by the Mann-Whitney *U* test when the *F*-value was significant. All statistical analyses considered *P* < 0.05 significant.

## ACKNOWLEDGEMENTS

This work was supported by funding from the National Institutes of Health K08HL128809 (B.T.C.) and in part by UPMC Children’s Hospital of Pittsburgh. Special thanks to Dr. Taylor Eddens for insightful discussions of this work.

## CONFLICT OF INTEREST

The authors declare no conflict of interest.

---

**Supplemental Figure 1. *Nr4a1*^*−/−*^ and WT mice have similar immune cell abundance following *K. pneumoniae* lung infection**

**(a)** The total amount of lung cells from WT (*n*=5) and *Nr4a1*^*−/−*^ (*n*=6) mice 24 hours post infection. The **(b)** total CD45^+^/CD11b^+^/CD11c^+^ (inflammatory macrophages), **(c)** CD45^+^/CD11c^+^/SiglecF^+^ (alveolar macrophages), and **(d)** CD45^+^/CD11b^+^/Ly6G^+^ (neutrophils) 24 hours after infection. Data are representative of two individual experiments.

**Supplemental Figure 2. *Nr4a1*^*−/−*^ and WT mice have similar cytokine expression after 24 hours of *K. pneumoniae* lung infection**

Transcript abundance for cytokines **(a)** *Il1β*, **(b)** *Tnfa*, and **(c)** *Il6*, as well as the neutrophil-specific gene **(d)** *Lcn2* from WT (*n*=6) and *Nr4a1*^*−/−*^ (*n*=8) whole lung. No significant differences were observed between genotypes 24 hours after *K. pneumoniae* pulmonary infection. All gene transcripts were normalized to *HPRT*.

**Supplemental Figure 3. FACS gating strategy for myeloid cells**

**Supplemental Figure 4. *Nr4a1*^*−/−*^ macrophages retain bactericidal function *in vitro***

**(a)** *K pneumoniae* burden from the supernatants of WT (*n*=5) and *Nr4a1*^*−/−*^ (*n*=4) bone marrow macrophages 30 minutes and 60 minutes after co-culture with *K. pneumoniae*. The gene transcript for **(b)** *Il1β*, **(c)** *Cxcl2*, and **(d)** *Il6* at 60 minutes exhibits no difference between WT and *Nr4a1*^*−/−*^ mice. All gene transcripts were normalized to *B2M*.

